# Granadaene Photobleaching Reduces the Virulence and Increases Antimicrobial Susceptibility of *Streptococcus agalactiae*

**DOI:** 10.1101/2020.03.31.019372

**Authors:** Sebastian Jusuf, Pu-Ting Dong, Jie Hui, Erlinda R. Ulloa, George Y. Liu, Ji-Xin Cheng

## Abstract

*Streptococcus agalactiae*, also known as Group B Streptococcus (GBS), is increasingly recognized as a major cause of soft tissue and invasive diseases in the elderly and diabetic populations. Antibiotics like penicillin are used with great frequency to treat these infections, although antimicrobial resistance is increasing among GBS strains and underlines a need for alternative methods not reliant on traditional antibiotics. GBS hemolysin/cytolysin and granadaene pigment are two major linked virulence factors that contribute to GBS pathogenicity. Here we show that photobleaching of the antioxidant granadaene renders the pathogen more susceptible to killing by mouse macrophages and to hydrogen peroxide killing. Photo-treatment also leads to loss of activity of the linked hemolysin/cytolysin although photobleaching disproportionally affected the activity of the two factors. Treatment with light also affected GBS membrane permeability and contribute to increased susceptibility to the cell membrane active antibiotic daptomycin and to penicillin. Overall our study demonstrates a dual effect of photobleaching on the virulence and antimicrobial susceptibility of GBS and suggests a novel approach for the treatment of GBS infection. Our findings further provide new insight on the relationship between GBS hemolysin and the granadaene pigment.

**Importance:** For elderly individuals or those with chronic underlying conditions (such as diabetes), skin infections caused by *Streptococcus agalactiae* represent a significant risk for the development of invasive disease. *S. agalactiae* strains are becoming increasingly resistant to antibiotics. By utilizing blue light to neutralize the granadaene pigment present in *S. agalactiae*, this paper presents a non-invasive and non-antibiotic reliant process capable of reducing GBS virulence while increasing the antimicrobial susceptibility of the bacterium. The differential effect of blue light on the linked GBS hemolysin/cytolysin and granadene pigment further provides new insight on the relationship between the two virulence factors. Overall photo-treatment represents a novel strategy for the treatment of *S. agalactiae* infections.

## Introduction

Gram-positive streptococci are responsible for a wide variety of medical conditions, ranging from more mild infections like strep throat to more serious infections such as meningitis, myositis, and necrotizing fasciitis (1–3). Among the many species within the *Streptococcus* genus, *Streptococcus agalactiae* (commonly referred to as group B *streptococcus* or GBS) is a beta-hemolyic, catalase negative facultative anaerobe that is best known for causing neonatal invasive diseases (4, 5). In recent years, there has been a greater appreciation for GBS as a leading cause of skin and soft tissue infections as well as invasive disease in the elderly and diabetic patients (6). While the risk of neonatal GBS infections has significantly decreased as a result of preventive intrapartum antibiotic prophylaxis (7, 8), the rate of GBS infection within non-pregnant adults has increased as evidence by a 2-fold rise in incidence of invasive GBS disease in adults between 1990 and 2007 (9). Compared to *S. pneumoniae* invasive diseases (42 per 100,000), a leading cause of mortality in the elderly population, the incidence of GBS invasive disease among elderly is remarkable at 25 per 100,000, based on active, population-based surveillance data from the ABCs Emerging Infections Program Network on cases of and deaths associated with infection due to several Gram-positive pathogens (6). As the rate of infection increases, treatment of GBS infection has become more reliant on the usage of antibiotics like penicillin V, ampicillin, and daptomycin (10, 11). However, even with optimal antibiotic treatment, management of skin infections can be difficult especially in patients suffering from diabetes (12). Compounding this therapeutic challenge, GBS has developed resistance to commonly used antibiotics like erythromycin and clindamycin, and has shown the capacity to develop resistance to beta-lactam antibiotics, threatening future use of current first-line drugs (13, 14). Because of these challenges, there is a growing need to develop novel GBS therapeutics that are less reliant on antibiotics.

One of the key reasons why GBS is highly infectious is its multitude of virulence factors that endow it with the ability to subvert suboptimal host defenses, allowing the bacterium to better adapt to its environment and cause severe infection (15). Some of these GBS virulence factors include the pore-forming toxin beta-hemolysin/cytolysin; the immune system evasion factor sialic acid capsular polysaccharide; as well as antimicrobial resistance factors like penicillin binding proteins and pili (16). Amongst all these different virulence factors, one of the most enigmatic is the ornithine rhamno-polyene pigment known as granadaene, which is localized within the plasma membrane of GBS (17). As the source of the characteristic red color of GBS, granadaene contains a linear chain of 12 conjugated double bonds, providing the pigment with an absorption spectrum nearly identical to most carotenoids, with absorption peaks present at 430, 460, 490, and 520 nm (18). This similarity in conjugated double bonds provides granadaene with antioxidant characteristics shared by traditional carotenoids, and provides it with the ability to quench and detoxify reactive oxygen species (ROS) like hydrogen peroxide (H_2_O_2_), singlet oxygen and superoxide (19). Previous studies have shown that pigment presence increases GBS resistance to ROS like H_2_O_2_ and superoxide by 5- to 10-fold compared to pigment deficient mutants, as well as significantly contributing to the survival within phagocytes by shielding the bacteria from oxidative damage (20). In addition to enhancing survival under oxidative stress, studies have indicated a link between the pigment expression and hemolytic activity of GBS, both of which are controlled by the *cyl* operon (21). While it has yet to be determined if the pigment and hemolysin are the same molecule, the findings that increased pigment production always correlates with increased hemolysin production and that isolated and stabilized granadaene pigment exhibits hemolytic activity, strongly indicate a close linkage between the two virulence determinants (22, 23).

Based on the multiple conjugated bonds located within the structure of granadaene, the granadaene pigment is highly likely to exhibit similar chemical reactivity to traditional carotenoids, such as the increased photosensitivity and photodegradation of the carotenoid structure in response to near UV and blue light exposure (24). Previous experiments have demonstrated that the staphyloxanthin (STX) carotenoid pigment within methicillin-resistant *S. aureus* (MRSA) is capable of undergoing photolysis when exposed to 460 nm light. This results in the formation of pores within the membrane of MRSA that not only increase the susceptibility of MRSA to reactive oxygen species, but also resensitize MRSA to conventional antibiotics the bacteria was previously resistant to (25, 26).

Considering the structural similarity between carotenoids and the granadaene pigment, we hypothesized that granadaene is capable of exhibiting similar photochemical reactions to blue light sources. In this paper, we present a novel approach for reducing virulence and increasing the antimicrobial susceptibility of infectious *S. agalactiae* through granadaene photobleaching. By degrading the pigment within the membrane, we significantly reduced the hemolytic activity of GBS and increased GBS susceptibility to oxidative stress, thereby reducing survival of the bacteria within macrophages. In addition, we demonstrated that photobleaching of GBS leads to improved effectiveness of antibiotics such as penicillin V and daptomycin. Based on its capacity for reducing bacterial virulence and increasing antimicrobial susceptibility, photobleaching potentially represents an effective adjunct therapeutic for the treatment of GBS skin and soft tissue infections.

## Results and Discussion

### Photobleaching of Granadaene Pigment and GBS

A previous study by Rosa-Fraile et al. established that the granadaene pigment expressed by *S. agalactiae* is a ornithine rhamno-polyene that contains a linear chain of 12 conjugated double bonds (**Figure 1a**) (17). Based on the protocol established by that study, we extracted the granadaene pigment from GBS (ATCC 12386) incubated in Strep B Carrot broth (**Figure 1b**). Using the extracted pigment solution, the absorbance profile of the pigment was measured and found to exhibit the absorbance peaks at 460 nm, 490 nm, and 520 nm (**Figure 1c**). In addition to these three explicit peaks, a small peak was observed at the 430 nm wavelength, which was also described in a previous report (18). Consistent with previous absorbance peak measurements of granadaene, it was established that the extracted solution contained the bacterial granadaene pigment. To determine overall photobleaching efficiency, we used pulsed light (60 J/cm^2^) with differing wavelengths and measured the greatest decrease in pigment peak. 430 nm light exhibited the greatest decrease in 490 nm pigment peak, with a peak decrease of 23.01% (**Figure S1**). Other wavelengths of light tested, such as 460 nm and 490 nm, were only found to decrease the pigment peak by only 10.86% and 5.1% respectively. Thus, it became apparent that exposure to 430 nm light was the most effective method for photobleaching the pigment. Exposure to 30 J/cm^2^, 60 J/cm^2^, and 120 J/cm^2^ of 430 nm blue light resulted in a 23.2%, 35.1%, and 39.6% reduction of the 490 nm absorbance peak respectively (**Figure 1d**). By the time the pigment solution had been exposed to 10 minutes of 430 nm light exposure, the entire peak profile of the pigment had completely disappeared, demonstrating the vulnerability of the granadaene pigment to 430 nm light exposure.

**Figure 1:**
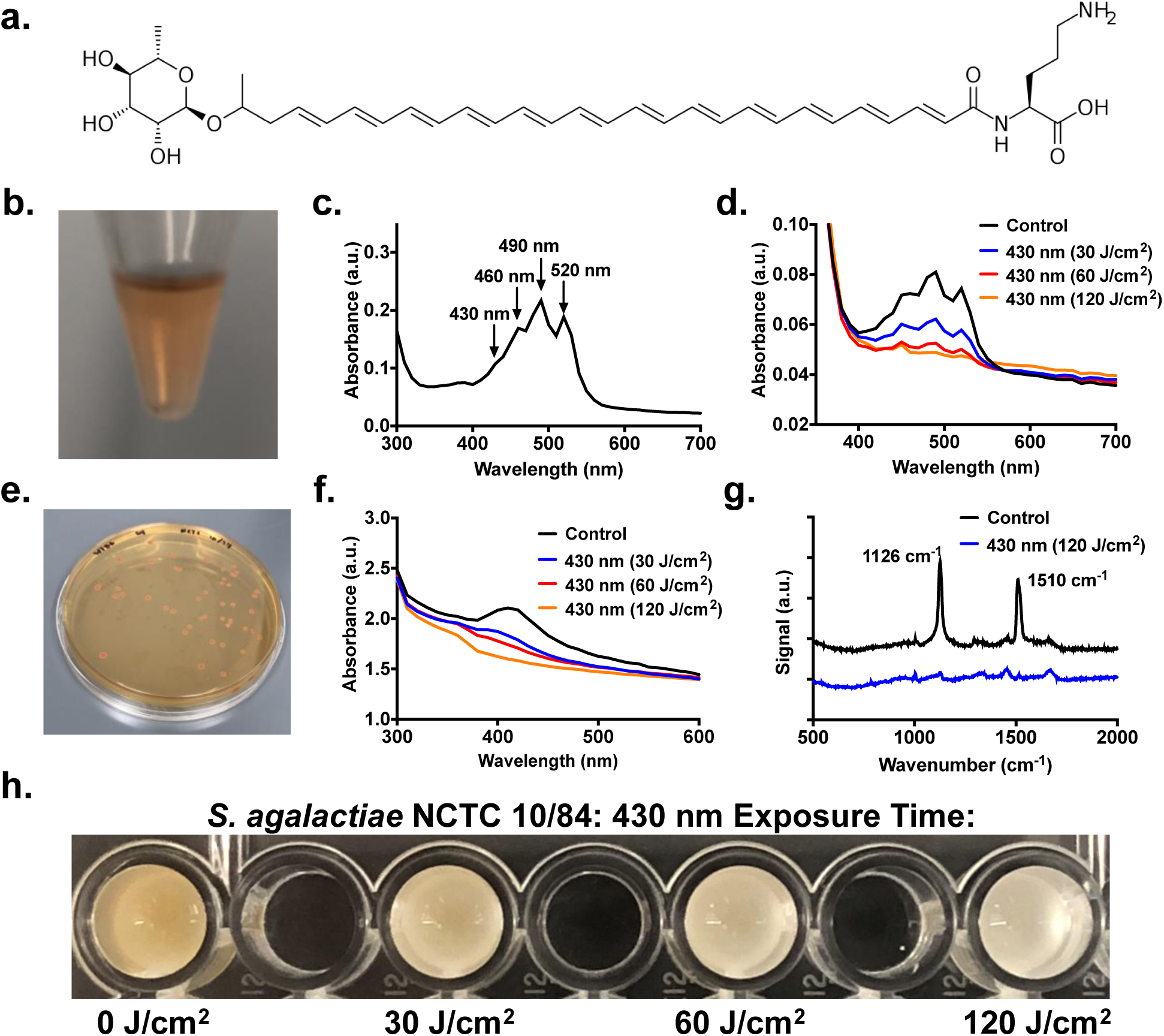
Extraction and photobleaching of granadaene pigment within *S. agalactiae*. (**a**) Chemical structure of granadaene. (**b**) Characteristic orange-red color of granadaene extracted from *S. agalactiae* (ATCC 12386) resuspended in DMSO with 0.1% Trifluoroacetic acid. (**c**) Absorption spectrum of extracted granadaene with characteristic peaks at 430 nm, 460 nm, 490 nm, and 520 nm. (**d**) Absorption spectrum of isolated granadaene following exposure to 0 (black), 30 (blue), 60 (red), and 120 J/cm^2^ (yellow) of nanosecond pulsed 430 nm blue light. Absorption peaks initially present at 490 nm disappears upon light exposure. (**e**) Granadaene-expressing *S. agalactiae* (NCTC 10/84) exhibits a strong orange-red color when grown on agar plates. (**f**) Absorption spectrum of concentrated *S. agalactiae* (NCTC 10/84) colonies following exposure to 0 (black), 30 (blue), 60 (red), and 120 J/cm^2^ (yellow) of 430 nm pulsed blue light. Absorption peak for pigment within bacteria appears to shift to 410 nm, which begins to disappear upon light exposure. (**g**) Raman spectroscopy of *S. agalactiae* (NCTC 10/84) following exposure to 120 J/cm^2^ of pulsed blue light. Characteristic peaks associated with carotenoid structures at 1126 cm^-1^ and 1510 cm^-1^ disappear following light exposure. (**h**) Photobleaching of granadaene results in visible change in color from orange to white in concentrated GBS. Bleaching increases with increasing light exposure.

Following photobleaching of the extracted pigment solution, GBS bacterial colonies exhibiting high pigment expression (NCTC 10/84) were cultured on agar plates and extensively used for photobleaching and subsequent studies. (**Figure 1e**) When these bacterial colonies were washed and concentrated in PBS, the absorbance profile of GBS colonies did not exhibit the expected absorption peaks found in the extracted pigment solution. Instead, the bacterial solution was found to express only a single absorption peak at 410-420 nm. While these results do not show the same three peak characteristics observed in the extracted pigment, a previous study by Rosa-Fraile et al. demonstrated that at high pH, the absorbance peaks of the granadaene were consolidated into a single peak at 420 nm (17). Thus, the 410 to 420 nm absorbance peak detected in the bacterial solution points to the presence of the pigment in the bacteria. Exposure of this bacterial solution to 530 J/cm^2^, 60 J/cm^2^, and 120 J/cm^2^ of 430 nm blue light was found to result in a 12.9%, 17.3%, and 24.1% reduction of the 410 nm absorbance peak respectively, thus exhibiting similar photobleaching capabilities to the extracted pigment (**Figure 1f**).

To better validate the presence of pigment within the cultured GBS colonies and photobleaching induced by light exposure, Raman spectroscopy was utilized to chemically detect the presence of pigment with GBS colonies (**Figure 1g**). Previous Raman studies have indicated that carotenoids exhibit intense spectral peaks within the 900 to 1600 cm^-1^ regions (27). In carotenoids, a strong peak at 1520 cm^-1^ (*v*_1_) peak is associated with the C=C stretching vibration, while a strong 1157 cm^-1^ (*v*_2_) peak is usually associated with C-C stretching. For our data, we observed that pigmented GBS exhibit significant Raman resonance peaks at the 1126 cm^-1^ and 1510 cm^-1^ wavenumbers. These results are consistent with the carotenoid Raman literature, as the 1510 cm^-1^ peak observed can be attributed to the C=C stretching vibration. While the 1126 cm^-1^ peak observed might appear inconsistent with the expected C-C stretching peak at 1160 cm^-^ 1, this can be explained by the lack of CH_3_ groups present on the granadaene pigment. Prior studies have indicated that the presence of CH_3_ groups on the carbon backbone interferes with C-C stretching, artificially increasing the resonance wavenumber. In polyacetylenic molecules that lack such CH_3_ groups, such as granadaene, the C-C peak can be expected to be generally 20 cm^-1^ lower than for carotenoids with the same chain length (27). Given this information, the observed resonance peak observed in granadaene at 1126 cm^-1^ could indeed be attributed to C-C stretching, confirming detection of the polyene structure of granadaene within the bacterial sample. With the strong presence of these polyene peaks within the bacterial sample established, it was determined that exposure to 120 J/cm^2^ was able to nearly eradicate the 1520 cm^-1^ and 1126 cm^-1^ peaks present within the sample, with a decrease in peak height of 90.2% and 88.4%, respectively. This reduction in Raman resonance peaks confirms the destruction of the pigment induced by exposure to 410 nm light. The inactivation of the pigment was further corroborated through visual means, as increasing exposure dosages of 410 nm light resulted in an increased loss of the distinct orange-red color associated with GBS, eventually reaching a blank white color after 120 J/cm^2^ of light exposure (**Figure 1h**).

### Granadaene Photobleaching Reduces GBS Hemolytic Activity

Having established that exposure to 430 nm light inactivates the granadaene pigment present within GBS, we next investigated the potential for photobleaching to inhibit the hemolytic capabilities of GBS. While several studies have demonstrated the close relationship between the β-hemolysin/cytolysin and the pigment within GBS, the specific mechanisms of the relationship are still not fully understood. However, pigment production and hemolytic activity are clearly and directly correlated with one another (22). Keeping this relationship in mind, the impact of pigment photobleaching on hemolytic activity was quantified for pigment producing GBS strains using the GBS NCTC 10/84 strain cultured under different growth conditions. For high pigment expressing GBS, GBS was cultured on tryptic soy agar plates and pigment production was allowed to accumulate. These colonies were then removed and allowed to grow in Todd Hewitt Broth for 48 hours (**Figure 2a**). Using a 1% blood solution, the hemolytic activity of non-exposed and 430 nm (120 J/cm^2^) exposed bacteria was calculated based on positive and negative controls utilizing 0.1% Triton X detergent and 1x PBS (**Figure 2b**). Based on these results, high pigment expressing GBS were found to express a hemolytic activity of 14.2%, while photobleached GBS expressed -0.1%, or 0% hemolytic activity (**Figure 2c**). In comparison, low pigment expressing GBS were prepared by culturing the GBS strain overnight in Todd Hewitt broth (**Figure 2d**). Following the same assay steps as the high pigment expressing GBS, the hemolytic activity of non-exposed and 430 nm (120 J/cm^2^) exposed bacteria was calculated (**Figure 2e**). The low pigment expressing GBS was found to express a hemolytic activity of 88.8%, while photobleached GBS expressed 48.3% hemolytic activity (**Figure 2f**).

**Figure 2:**
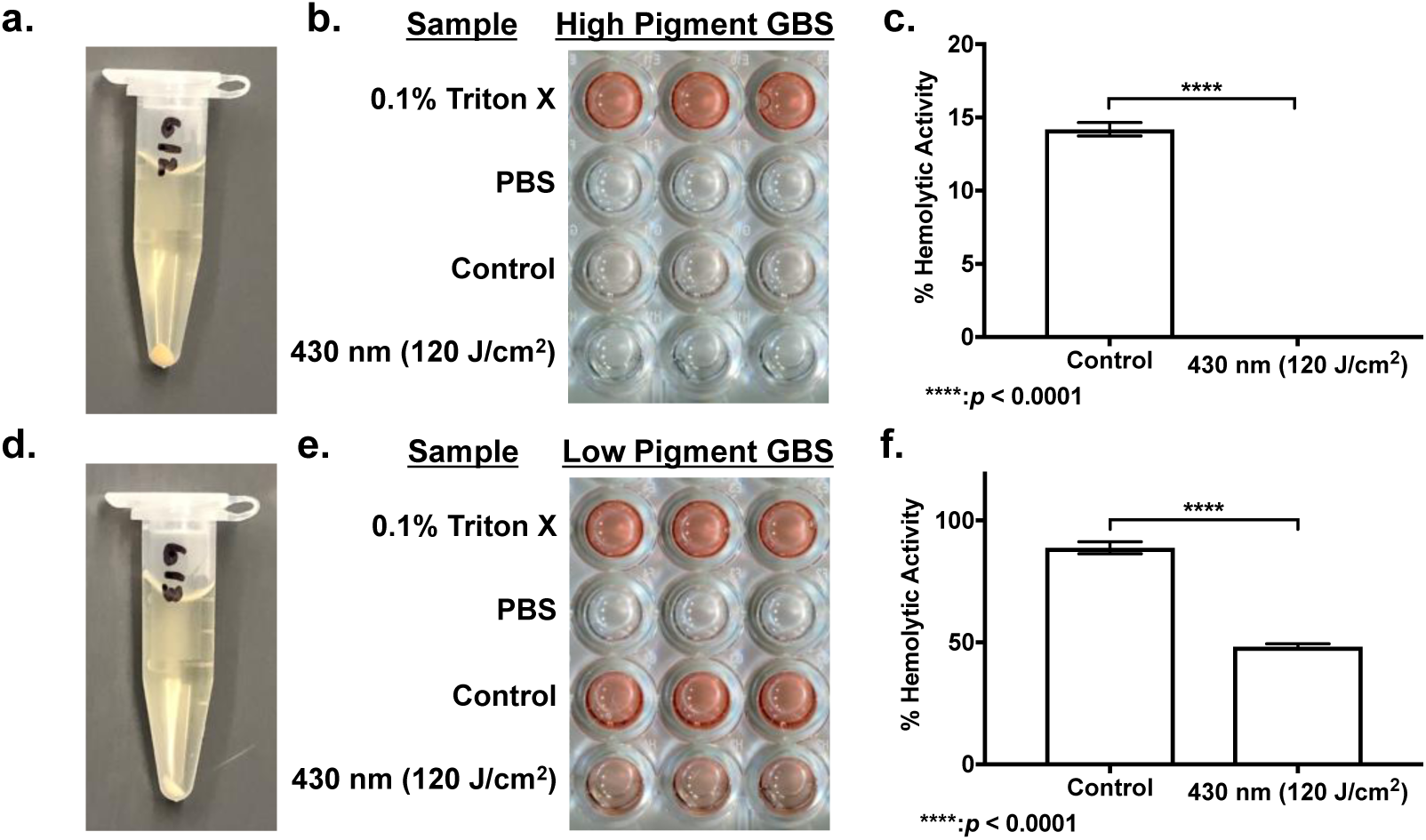
Granadaene photobleaching reduces Hemolytic activity in *S. agalactiae*. (a) Centrifuged pellet of agar plate grown GBS exhibits distinct orange pigmentation. (b) Hemolytic assay of high pigment expressing GBS with photobleaching alongside positive and negative controls. (c) Percent hemolytic activity of non-exposed and blue light exposed high pigment expressing GBS. (d) Centrifuged pellet of overnight broth cultured GBS exhibits significantly lower pigmentation. (b) Hemolytic assay of low pigment expressing GBS with photobleaching alongside positive and negative controls. (c) Percent hemolytic activity of non-exposed and blue light exposed low pigment expressing GBS. ****: *p* < 0.0001.

For both GBS preparations tested, a reduction in hemolytic activity was observed immediately following photobleaching. For the high-pigment expressing GBS, photobleaching completely disabled all hemolytic activity, whereas photobleaching reduced hemolytic activity by 40.5% in the low pigment expressing GBS. For the high pigment expressing GBS, the initial hemolytic activity observed was found to be rather low at 14.2%. Previous routine observations of GBS have found that as GBS progresses further into stationary phase, the hemolytic activity decreases while the pigmentation appears to increase. Regardless, both samples were found to experience significant reductions in hemolytic activity, suggesting that the presence of the pigment does indeed play a significant role in the hemolytic activity of GBS, and destruction of the pigment through photobleaching is capable of significantly inhibiting the hemolytic capabilities of the GBS bacterium. However, a notable finding was how the hemolytic activity of the low pigment expressing GBS was elevated at 88.8% at the start and how photobleaching only reduced activity by approximately 40%. These results indicate that photobleaching is capable of significantly reducing the hemolytic activity of GBS, as well as suggesting that an overlap model exists between granadaene and the hemolysin regarding the overall hemolytic activity of GBS. While granadaene does contribute to the total hemolytic activity of GBS, the residual hemolytic activity measured after photobleaching indicates that the GBS hemolysin/cytolysin primarily responsible for hemolytic activity is not identical in structure to the pigment, although they are like closely related or overlapping in structure.

### Granadaene Photobleaching Sensitizes GBS to H_2_O_2_ and Macrophage Attack

Previous work by our group have demonstrated that the granadaene pigment production in GBS confers resistance to reactive oxygen species (ROS) (20). With this in mind, it was found that application of 60 J/cm^2^ of 430 nm light and 12 mM of H_2_O_2_ resulted in the complete eradication of pigment expressing wild type (WT) GBS (**Figure 3a**). Light exposure alone was found to have no significant effect on the survival of WT GBS, while the addition of 12 mM of H_2_O_2_ resulted in only a 92.5% reduction in CFU/mL. In contrast, when non-pigmented Δ*CylE* GBS mutant cultures were tested, the combination of light and 16 mM of H_2_O_2_ resulted in only a 93.4% CFU reduction from the Δ*CylE* GBS treated with only 16 mM of H_2_O_2_ (**Figure 3b**). When the effectiveness of H_2_O_2_ in killing either the pigmented or non-pigmented GBS was compared, it was observed that, for H_2_O_2_ only samples, WT GBS CFU were reduced by only one log (92.5%) CFU while Δ*CylE* GBS CFU was reduced by 2 logs (98.4%), indicating that Δ*CylE* GBS was generally more susceptible to ROS consistent with previous studies. The complete eradication of WT GBS following photobleaching indicates that photobleaching is capable of significantly reducing antioxidant activity present within GBS, and by doing so, removing the ability for GBS to resist oxidative stress, a phenomena that is not observed to the same extent in Δ*CylE* GBS.

**Figure 3:**
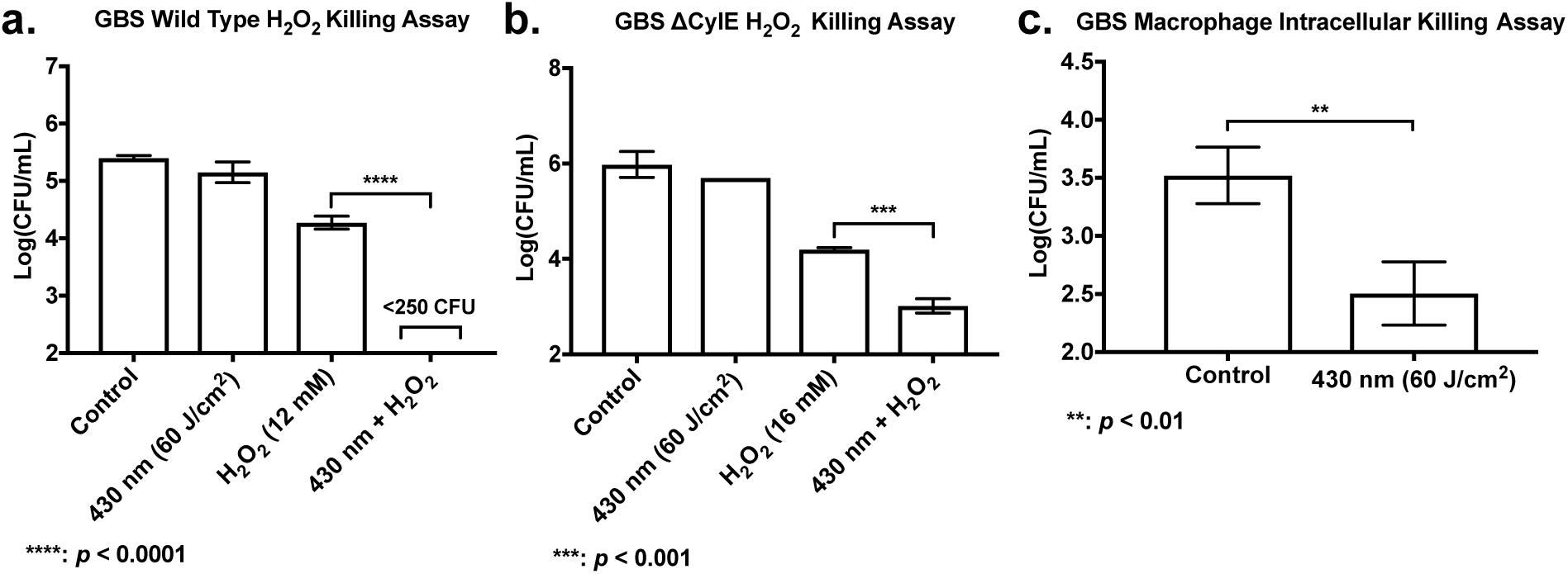
H_2_O_2_ killing assays of wild type and mutant GBS alongside WT GBS macrophage intracellular killing assay. (a) Pigment expressing GBS exposed to 60 J/cm^2^ of 430 nm nanosecond pulsed light and incubated with 12 mM of H_2_O_2_ for 1 hour. Combination of 430 nm light exposure and H_2_O_2_ resulted in complete eradication of GBS. (b) Pigment deficient Δ*CylE* GBS exposed to 60 J/cm^2^ of pulsed light and incubated with 16 mM of H_2_O_2_ for 1 hour. Combination of 430 nm light exposure and H_2_O_2_ resulted in only 1 log improvement in antimicrobial activity. (c) CFU count of pigment expressing GBS engulfed by RAW 264.7 murine macrophages and exposed to 60 J/cm^2^ of 430 nm light. Survival CFU of GBS was assessed following 8 hours of incubation in DMEM. ****: *p* < 0.0001; ***: *p* < 0.001; **: *p* < 0.01.

Since photobleaching using a 430nm light source reduces the capacity of GBS to resist ROS killing, we next explored the survival of photobleached WT GBS within murine macrophages (**Figure 3c**). With photobleaching, GBS survival within macrophages was one log (90.4%) lower compared to non-photobleached bacteria after 8 hours. These results were consistent with the results of the H_2_O_2_ GBS killing assay and demonstrate that photobleaching of GBS granadaene significantly reduces the ability of GBS to survive host mediated oxidant stress.

### Granadaene Photobleaching Sensitizes GBS to Antibiotics

Since granadaene pigment is localized within the membrane of *S. agalactaie*, we sought next to quantify the impact of photobleaching on bacterial membrane permeability using SYTOX Green (**Figure 4a**). The results demonstrate that exposure to 430 nm light results in a near instant increase in the SYTOX Green fluorescence, with a 35.9% increase in measured fluorescence following 30 J/cm^2^ dosages from the control at time point t = 0 minutes. By 60 minutes, the fluorescence of GBS exposed to 30 J/cm^2^ of light increased by an average of 44.2% respectively. As a fluorescent dye with a high affinity for nucleic acids, changes in SYTOX Green fluorescence is an ideal measurement to quantify alteration in membrane permeability of bacteria (28). Since SYTOX is normally membrane impermeable, the dye only attaches and fluoresces to nucleic acids found in membrane compromised bacteria. Thus, the significant increases in measured fluorescence for 430 nm is indicative of the membrane disrupting effects of granadaene photobleaching.

**Figure 4:**
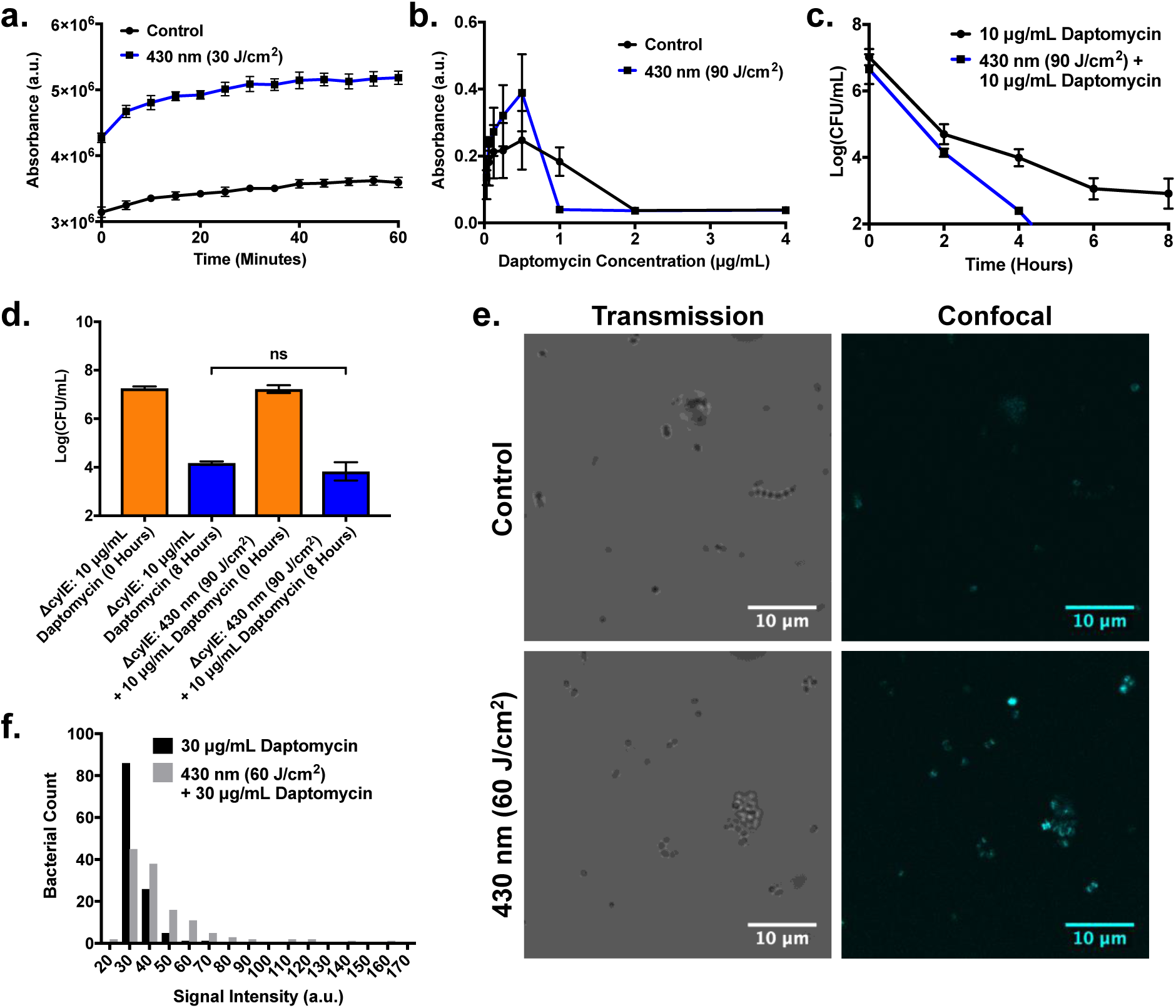
Granadaene photobleaching facilitates cellular uptake and antimicrobial activity of daptomycin. (a) Membrane permeability assay of GBS exposed to 430 nm nanosecond pulsed light utilizing SYTOX Green. Instantaneous increase in SYTOX Green fluorescence following light exposure indicates immediate membrane permeabilization. (b) Minimum Inhibitory Concentration (MIC) of daptomycin on GBS photobleached with blue light decreases 2-fold from 2 μg/mL to 1 μg/mL. (c) CFU count of time killing assay of GBS exposed to 90 J/cm^2^ of pulsed blue light. GBS exhibits faster response to daptomycin following initial light treatment. (d) CFU count comparison of pigment deficient Δ*CylE* GBS exposed to 90 J/cm^2^ of pulsed blue light and treated with 10 μg/mL for 8 hours. Light exposed Δ*CylE* GBS lacks the same daptomycin enhancement observed in the pigmented strain. (e) Confocal imaging of light exposed GBS treated with 30 μg/mL of daptoymcin-BODIPY for 30 minutes. The light treated GBS exhibits greater BODIPY fluorescence compared to the control, indicating higher daptomycin intake. (f) Histogram of signal intensities of GBS bacteria treated with 30 μg/mL of daptoymcin-BODIPY for 30 minutes. Light treated GBS exhibits a greater number of bacteria with higher signal intensities compared to the control.

Since photobleaching increased membrane permeability of GBS, we next studied the impact of increased membrane permeability on the effectiveness of various antibiotics such as daptomycin, penicillin V, and ampicillin. Following exposure to 90 J/cm^2^ of 430 nm light GBS showed a twofold decrease in daptomycin minimal inhibitory concentration (MIC), from 2 μg/mL to 1 μg/mL, (**Figure 4b**). A time kill assay of GBS treated with 10 μg/mL of daptomycin and 90 J/cm^2^ of 430 nm light demonstrated an additional 2 log reduction in CFU with light therapy and daptomycin compared to daptomycin alone after 4 hours in a TSB solution (**Figure 4c**). These results indicate that granadaene photobleaching can increase the effectiveness of daptomycin in clearance of GBS. To query the role of the granadaene pigment in the improved daptomycin performance, non-pigment expressing Δ*CylE* GBS was evaluated. In contrast to wild type GBS, survival of Δ*CylE* GBS was not statistically significant with daptomycin only and light plus daptomycin treatment (**Figure 4d**). The reduced effectiveness of photobleaching for the pigment-deficient GBS suggests that the removal of the pigment increases the performance of daptomycin. As a lipopeptide antibiotic, daptomycin induces damage on Gram-positive bacteria through its ability to disrupt the cell membrane of bacteria (11, 29). This mechanism of action is reliant on the binding and aggregation of daptomycin within the cell membrane, altering the curvature of the membrane and generating ion leak holes within the membrane that rapidly depolarize the bacterial environment. This alteration in membrane potential causes the inhibition of protein synthesis, leading to cell death (30). Given the membrane disrupting capabilities of granadaene photobleaching, it is possible that the membrane insertion and aggregation capabilities of daptomycin are enhanced given following photobleaching. To corroborate this hypothesis, light exposed and non-light exposed GBS was treated with BODIPY dye conjugated daptomycin and imaged using a confocal microscope (**Figure 4e**). Based on the images taken, the thresholding and particle tracking capabilities of ImageJ were used to obtain the average intensities of individual cells within the exposed and non-exposed GBS groups, all of which were charted onto a histogram (**Figure 4f**). Based on the histogram, the average cell intensity of the daptomycin only treated GBS was 33.12 ± 6.6, while the average intensity of the daptomycin plus light treated GBS was 46.11 ± 22.71. While the higher average intensity of the daptomycin plus light treated GBS indicates higher average daptomycin intake of the cells, the significantly higher standard deviation observed can be attributed to a combination of individual high intensity (120+) GBS cells as well. However, even with these outliers, a greater number of light treated GBS exhibit higher average intensities than the non-treated GBS, with 47.66% of photobleached GBS exhibiting average signal intensities above 40, compared with the 8.4% observed for control GBS. Given these trends, it is clear that photobleached GBS are capable of improved daptomycin aggregation based on increased fluorescence, allowing for improved efficiency and performance of daptomycin in 430 nm light exposed GBS.

Because daptomycin has an unique membrane-dependent mechanism of action, other antibiotics commonly used to treat GBS infections, such as ampicillin and penicillin V, were investigated to examine the potential effect of granadaene photobleaching on their effectiveness (31). For ampicillin, a time kill assay on both pigment-expressing GBS (**Figure 5a**) and the isogenic pigment deficient Δ*CylE* GBS (**Figure 5b**) showed that light exposure had no statistically significant impact on the effectiveness of ampicillin. However, for penicillin V treatment, 60 J/cm^2^ of 430 nm exposure augmented CFU reduction by 79.7% for the pigment-expressing GBS (**Figure 5c**) and by 67.2% for the pigment deficient Δ*CylE* GBS after 6 hours of incubation (**Figure 5d**). Interestingly, despite both ampicillin and penicillin V being part of the same subclass of beta lactam antibiotics, only penicillin V was found to exhibit significant improvement in performance (32). Given that both antibiotics induce antibacterial activity by inhibiting the synthesis of the peptidoglycan layer in bacteria by binding to penicillin binding proteins responsible for catalyzing peptidoglycan crosslinks, the different responses could be attributed to differences in the antibiotic chemical structure (33). Ampicillin is known to contain an additional amino group in their side chain that naturally increases the hydrophilic nature of the compound. While this increased hydrophilicity is generally understood to help ampicillin pass through porins present within the outer membrane of Gram-negative bacteria, this property may contribute to its improved access to the penicillin binding proteins present within the membrane and interior of the cell, therefore reducing the advantages induced by photobleached triggered bacterial membrane permeability (32, 34). Irrespective of the ineffectiveness of ampicillin, the improvement in penicillin V effectiveness indicates that photobleaching of the granadaene pigment is capable of improving, albeit modestly, penicillin V access the penicillin binding proteins present within the cell membrane and interior of the cell. This improvement in penicillin V efficacy may also be due to the increased susceptibility of photobleached pigment producing GBS to ROS. Previous studies have indicated that beta-lactam antibiotics such as penicillin are capable of inducing hydroxyl radical formation, and the removal of the granadaene antioxidant through photobleaching could slightly improve the antimicrobial activity of antibiotic induced ROS, contributing to the improved activity (35, 36).

**Figure 5:**
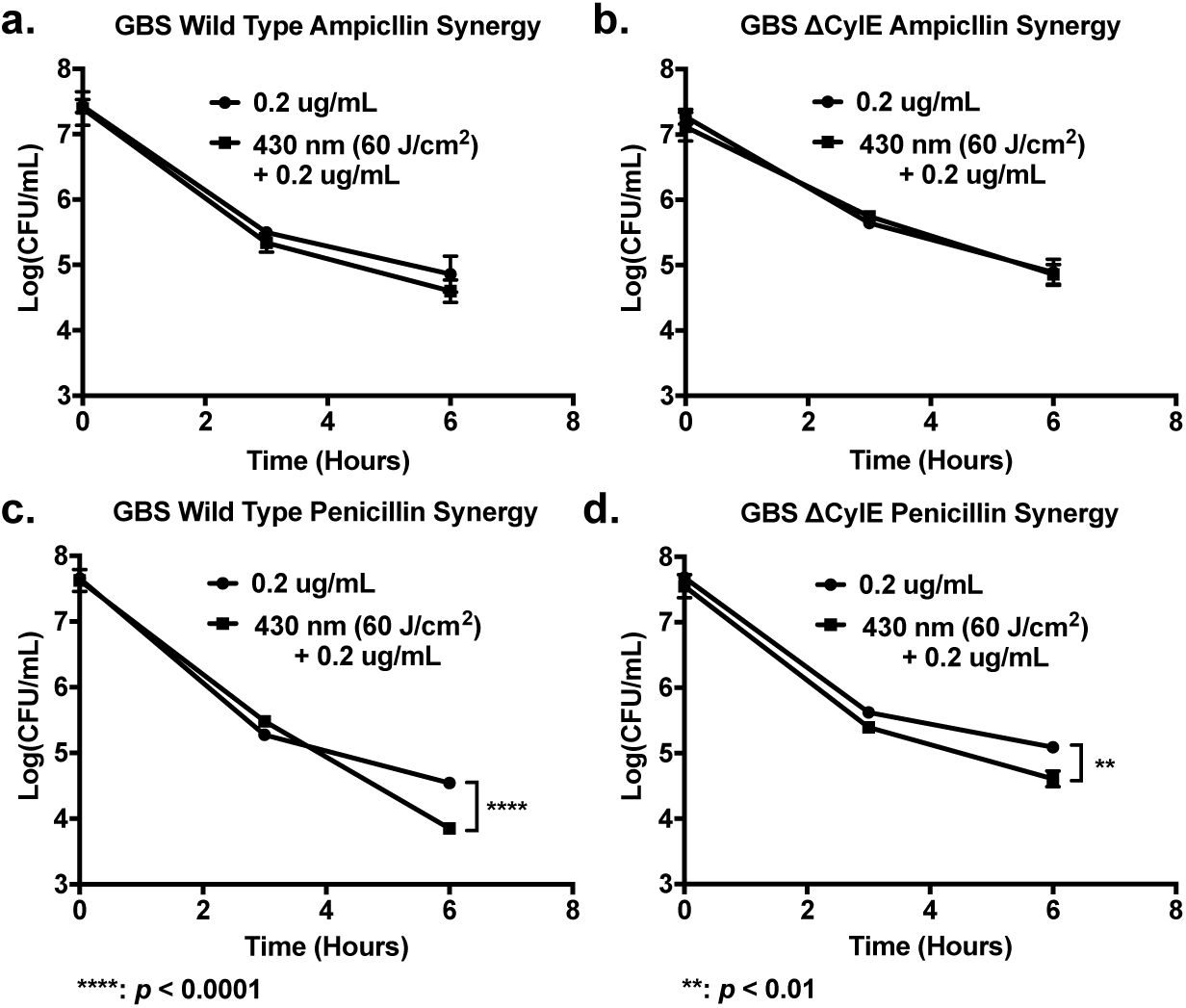
Granadaene photobleaching potentiates beta-lactam antibiotic activity on GBS. (a) CFU Time kill assay of GBS exposed to 430 nm pulsed light and treated with 0.2 μg/mL of ampicillin. No significant changes in ampicillin performance observed. (b) CFU Time kill assay of Δ*CylE* GBS exposed to 430 nm pulsed light and treated with 0.2 μg/mL of ampicillin. No significant changes in ampicillin performance observed. (c) CFU Time kill assay of GBS exposed to 430 nm pulsed light and treated with 0.2 μg/mL of penicillin V. Slight enhancement of penicillin V activity observed after 6 hours of incubation. (d) CFU Time kill assay of Δ*CylE* GBS exposed to 430 nm pulsed light and treated with 0.2 μg/mL of penicillin V. Minor enhancement of penicillin V activity observed after 6 hours of incubation, although not as large as the enhancement observed in the pigment expressing GBS. ****: *p* < 0.0001; **: *p* < 0.01.

Given the increasing development of reduced penicillin susceptibility in GBS, it is possible that photobleaching may provide an avenue to reduce potential penicillin resistance in GBS if it were to become widespread within the population (37, 38). However, it is clear with the current results that the pigment photobleaching process is more effective with antimicrobials like daptomycin that are dependent on membrane insertion and disruption.

One interesting finding that was observed with both daptomycin and penicillin V treatment of GBS was the modest increase in antibiotic efficacy when combined with light treatment, even when applied to the pigment-deficient Δ*CylE* GBS. To further probe this effect, we examined the impact of 430 nm light exposure on Δ*CylE* GBS. When the absorbance spectrum of Δ*CylE* GBS was examined, no significant peaks were observed in the control sample and light exposure had no effect on the absorbance spectrum (**Figure S2a**). However, when analyzed using the Raman spectroscopy, significant peaks were found to be present at the 1126 cm^-1^ and 1512 cm^-1^ (**Figure S2b**). These peaks were found to decrease by 66% when exposed to 60 J/cm^2^ of 430 nm light. In addition, membrane permeability studies found that, under 30 J/cm^2^ of 430 nm light, Δ*CylE* GBS exhibited a 19.8% increase in measured fluorescence immediately following bleaching at time point t = 0 minutes (**Figure S2c**). By 60 minutes, the fluorescence of GBS exposed to 30 J/cm^2^ of light increased by an average of 62.8% respectively. Based on published data, one potential explanation for these results can be attributed how the granadaene pigment is manufactured under the *cyl* operon (21). In the *cyl* metabolic pathway, production of the granadaene pigment begins through the creation of a 12-carbon double conjugated fatty acid through 12 cycles of malonyl-ACP additions to a growing crotonyl-AcpC chain (**Figure S2d**). While removal of the *CylE* gene abrogated pigment production in Δ*CylE* GBS, the generation of fatty acid chains through the *cyl* operon could still occur without the CylE protein, which is an acyl CoA acyltransferase responsible for the attachment of the L-ornithine to the end of the fatty acid chain. Given this functionality, it is likely that double conjugated fatty acid chains are still generated within the GBS, even if these chains do not lead to pigment formation. The increased membrane fluidity observed following light exposure could also be an indication that these fatty acids can still be incorporated into the plasma membrane of the cell, with the exposure to 430 nm disrupting the membrane integrity and permeability of the GBS. These results indicate that while the pigment itself is not necessary to disrupt the membrane integrity of GBS, the presence of the pigment allows for the photobleaching process to reduce the antioxidant activity and antibiotic susceptibility of GBS to a greater extent, as observed with both hydrogen peroxide and daptomycin.

Overall, this paper presents a non-antibiotic reliant method of reducing the hemolytic and antioxidant activity of GBS, thereby reducing the potential virulence and infectivity of the bacteria as well as improving the ability for macrophages to properly kill phagocytosized GBS. In addition, 430 nm exposure was also shown to disrupt membrane permeability of GBS and improve the activity of antibiotics like daptomycin. Based on these results, phototherapy could potentially enhance current treatment of *S. agalactiae* skin and soft tissue infection in adult patients, which would reduce the risk of life threatening invasive infections such as bacteremia and bone/joint infections (10, 39, 40). The reduced survivability of GBS within macrophages following photobleaching could be particularly useful for treatment of immunocompromised patients, such as elderly or diabetic individuals, with GBS infection (6, 41).

While granadaene photobleaching has the dual effect of reducing virulence and increasing antimicrobial susceptibility, other potential applications of photo treatment exist. While photobleaching induced only modest improvement in beta lactam antibiotic activity, it is conceivable that photobleaching would have a more pronounced effect on GBS strains that are moderately resistant to antibiotics. Therefore, future photobleaching studies will focus on the potential to improve penicillin treatment in *S. agalactiae* strains that are moderately resistant to penicillin. Additional studies will address if pigment photobleaching could resensitize resistant *S. agalactiae* strains to conventional antibiotics such as erythromycin and clindamycin (42). Overall, *S. agalactiae* granadaene photobleaching presents a method of reducing virulence and increasing antimicrobial susceptibility, creating a non-invasive way to improve upon current treatment of hard to treat GBS infections.

## Materials and Methods

### Source of blue light

Blue light was administered through a short-pulsed laser (OPOTEK). The laser is tunable from 410 nm to 2200 nm at a repetition rate of 20 Hz and a pulse width of ∼5 ns. At 430 nm, the system has a maximum pulse energy of 9 mJ. While the standard beam diameter of the system is ∼2 mm, a collimator attached to an optical fiber was used to expand the diameter of the beam to 10 mm. With these parameters, the pulsed laser offers a power output of ∼100 mW/cm^2^.

### Bacteria

*S. agalactiae* Lehmann and Neumann (ATCC 12386) were purchased from the American Type Culture Collection, and GBS grown in a Group B *Streptococcus* Carrot Broth was used for initial granadaene pigment extraction. For bacterial susceptibility and hemolytic assays, a highly pigment *S. agalactiae* strain (NCTC 10/84) and its non-pigmented isogenic Δ*CylE* mutant were used (provided by Victor Nizet, *Pritzlaff C Molecular Microbiology 2001*)

### Pigment Extraction

Extraction of the granadaene pigment was performed based on a modified extraction protocol published by Rosa-Fraile et al (17). To summarize, GBS (ATCC 12386) was grown in New Granada Media/Strep B Carrot Broth (Z40, Hardy Diagnostics) at 37°C for 48 to 72 hours until the broth turned a rich orange-red color. Bacterial suspension was centrifuged and washed three times with distilled water and two times with DMSO. Following washing, the bacterial pellet was resuspended in DMSO with 0.1% Trifluoroacetic acid (TFA) overnight. The next day, the suspension was centrifuged and the supernatant containing the extracted pigment was collected.

### Photobleaching and Absorbance/Raman Measurements

Photobleaching and absorbance measurements of extracted pigment and bacteria samples were performed by placing 90 µL of extracted pigment suspended in DMSO/0.1% TFA or concentrated GBS bacteria suspended in PBS in a 96 well plate. Once plated, the samples were exposed to 430 nm light for different time intervals (0, 5, 10, 20 minutes). The absorbance spectrum of each sample was measured through a plate reader spectrometer.

For Raman measurements, a Raman spectroscopy system (1221, LABRAM HR EVO, Horiba) with a 40x Olympus objective and an excitation wavelength of 532 nm was used to detect changes in pigment presence within GBS. Images were acquired under a 30 second acquisition time with a laser ND filter of 10%. Pigment-expressing GBS was concentrated in 50 µL of PBS and a 2 µL droplet was aliquoted and sandwiched between two cover slides (48393-230, VWR International). Changes in pigment presence were determined by measuring the change in Raman peak amplitudes associated with conjugated double bonds located at the 1126 cm^-1^ and 1510 cm^-1^ before and after light treatment. For Raman measurements collected with high background peaks, Raman Spectrum Baseline Removal Matlab Code was utilized to better isolate Raman peaks (43, 44).

### Hemolytic Activity Assay

Hemolytic activity of an agar plate- or broth-cultured NCTC 10/84 GBS was measured. The bacteria were washed and resuspended in 1 mL of 1x PBS. A 15 µL aliquot of bacterial suspension was exposed to 20 minutes of 430 nm light. Following the light exposure, the suspension was diluted with 35 µL of PBS and then further combined with 200 µL of 1% fresh human blood solution. Blood samples were acquired through the Boston Children’s Hospital Blood Donor Center. For positive controls, 200 µL of blood solution was mixed with 50 µL of 0.1% Triton X. For negative controls, 200 µL of blood solution was mixed with 50 µL of 1x PBS. Blood samples were cultured at 37°C for 2 hours, after which the samples were centrifuged and the supernatant were collected. The absorbance of the supernatant was measured at 420 nm. Percent hemolytic activity was determined by subtracting the GBS supernatant absorbance by the negative control absorbance, followed by dividing it all by the difference between the positive and negative control absorbance measurements.

### In Vitro Synergy Assessment between Photobleaching and Reactive Oxygen Species in GBS

*S. agalactiae* NCTC 10/84 strains (WT and non-pigment expressing Δ*CylE* mutant) were streaked and cultured on trypticase soy agar plates in a static 37°C incubator for 24 to 48 hours to maximize colony size. Once pigment-expressing colonies were formed, colonies were picked with a bacterial loop and then wash and resuspended in 1x PBS. A 10 µL aliquot of this bacterial suspension was placed onto a cover slide and exposed to 100 mW 430 nm pulsed light for various time intervals. Following exposure, the droplets were removed from the cover slide and mixed with 990 µL of PBS supplemented with different concentrations of hydrogen peroxide. These tubes were cultured for 1 hour. Once culturing was complete, serial 10-fold dilutions were performed using a 96 well plate. These dilutions were plated on tryptic soy agar plates in triplicate to quantify the number of surviving GBS colonies. Plates were incubated at 37°C overnight before counting CFU/mL.

### Intracellular GBS Macrophage Killing Assay

The intracellular GBS macrophage killing assay was adapted from a previously published protocol (20). To summarize, 10^5^ cells of RAW 264.7 murine macrophages (ATCC® TIB-71™) suspended in Dulbecco’s Modified Eagle Medium (DMEM) supplemented with 10% fetal bovine serum (FBS) were seeded into 96 well plates and allowed to adhere to the well surface overnight. Following adhesion, DMEM media was removed. To achieve a multiplicity of infection of 100, the OD_600_ of pigmented GBS resuspended in PBS was determined. Based on the OD_600_, a 20 µL droplet of GBS suspension containing 10^7^ CFU was added to each well alongside an addition 80 µL of DMEM. Bacteria were incubated with macrophages for 1 hour within a 37°C incubator, after which the bacterial media was removed and washed twice with pre-warmed clear DMEM (DMEM containing no phenol red). Once washing was complete, 100 µL of pre-warmed clear DMEM was added to each well. Each well was exposed to 10 minutes of 430 nm light. After light exposure, wells were washed once again with DMEM. To kill off any remaining extracellular GBS within the wells, each well as filled with 100 µL of DMEM with 10% FBS supplemented with 5 µg/mL of penicillin V and 10 µg/mL of gentamicin. The macrophages were then incubated for an additional 8 hours at 37°C. Following incubation, wells were washed 3 times with PBS and treated with 0.1% Triton X for 5 minutes to lyse macrophages and release surviving GBS. Serial dilutions were then performed and plated on agar plates to count the CFU/mL.

### Membrane Permeability Assay

*S. agalactiae* NCTC 10/84 colonies cultured on agar plates were removed, washed, and resuspended in in 100 µL of 1x PBS. A 10 µL aliquot of suspension was placed on a cover slide and exposed to 100 mW 430 nm light for 5 to 10 minutes. Following exposure, the exposed droplet was combined with 980 µL of sterile water and 10 µL of 0.5 mM SYTOX Green (S7020, Thermo Fisher Scientific) solution before being added to a 96 well plate (200 µL solution per well). Using a plate reader, the fluorescence of each well was measured over the course of 1 hour at 5-minute intervals. The excitation wavelength was set at 488 nm while the emission wavelength was set at 525 nm. Increasing emission intensity was indicative of greater membrane permeability.

### In Vitro Synergy Assessment between Photobleaching and Antibiotics

For initial minimum inhibitory concentration determinations, agar plate-grown *S. agalactiae* NCTC 10/84 colonies were washed and resuspended in 1x PBS. A 5 µL aliquot of this bacterial suspension was placed onto a cover slide and exposed to 100 mW 430 nm pulsed light for specific time intervals. Following exposure, the droplets were removed from the cover slide and mixed with 5 mL of Todd Hewitt Broth. (T1438-100G, Sigma Aldrich). 100 µL of GBS containing broth was added to a 96 well plate, after which an additional 100 µL of GBS broth was added to the first row of the plate alongside an initial starting concentration of antibiotics daptomycin (103060-53-3, Acros Organics), penicillin V (1504489, Sigma Aldrich), ampicillin (A9518, Sigma Aldrich). 2-fold serial dilutions were performed and the plate was allowed to incubate overnight. Following overnight culture, the plate was removed and measured through both visual means and through a plate reader to determine the minimum inhibitory concentration of different antibiotics. For daptomycin studies, initial broth cultures were supplemented with 50 µg/mL of CaCl_2_.

To determine potential synergy, *S. agalactiae* NCTC 10/84 colonies were removed from agar plates and then washed and resuspended in 1x PBS. A 10 µL aliquot of this bacterial suspension was placed onto a cover slide and exposed to 430 nm pulsed light for various time intervals. Following exposure, the droplets were removed from the cover slide and mixed with 990 µL of PBS supplemented with differing concentrations of daptomycin, penicillin V, and ampicillin. For daptomycin studies, cultures were supplemented with 50 µg/mL of CaCl_2_. These culture tubes were incubated at 37°C and CFU was measured through serial dilutions at various time intervals.

### Confocal Imaging and Tracking of Daptomycin Membrane Integration

To visualize the integration of daptomycin into the membrane of GBS using confocal imaging, daptomycin was conjugated to the fluorescent marker BODIPY STP-ester. To prepare this drug-dye combination, 10 mg of daptomycin was dissolved in 1 mL of 0.1 M NaHCO_3_ solution. 100 µL of 1 mg/mL BODIPY STP-ester (B10006, Thermo Fisher Scientific) was added to the daptomycin solution drop by drop and then stirred for 1 hour at room temperature. Following this, the solution underwent dialysis utilizing 0.1 M NaHCO_3_ solution overnight. Once dialysis was complete, the drug-dye complex was lyophilized.

In order to prepare samples for imaging, pigment expressing GBS colonies were removed from agar plates, centrifuged, washed, and resuspended in 100 µL of PBS. A 10 µL aliquot of GBS was placed on a cover slide and exposed to 10 minutes of 430 nm light. Following exposure, the 10 µL aliquot was combined with 970 µL of Todd Hewitt Broth, 10 µL of 5 mg/mL CaCl_2_, and 10 µL of 3 mg/mL Daptomycin-BODIPY. The GBS culture was incubated for 30 minutes at 37°C, after which the culture was washed with PBS and fixed in 50 µL of 10% formalin. A 2.5 µL aliquot of GBS fixed in formalin was sandwiched between a Poly-L lysine coated glass slide and a cover slide.

For confocal imaging, a FV3000 Confocal Laser Scanning Microscope (Olympus) was used to image and detect the presence of daptomycin within the membrane of GBS. Confocal images were acquired under an excitation wavelength of 488 nm and an emission detection wavelength range of 510 nm to 560 nm. Images were taken under the following confocal settings: 100x Objective, 2.85x Zoom, a PMT Voltage of 600 V, and a laser ND Filter of 10%. Images were processed through ImageJ utilizing thresholding and particle analysis in order to obtain the average intensities of individual cells.

### Statistical Analysis

All comparative data was analyzed utilizing a student t-test to obtain significance between data sets. All figures and analysis were performed using ImageJ or GraphPad PRISM version 7.

## Acknowledgements

We thank the Blood Donor Center at the Boston Children’s Hospital for providing blood samples.

Funding for this project was supported by the Boston University Start Up Fund.

